# The spindle assembly checkpoint functions during early development in non-chordate embryos

**DOI:** 10.1101/582759

**Authors:** Janet Chenevert, Marianne Roca, Lydia Besnardeau, Antonella Ruggiero, Dalileh Nabi, Alex McDougall, Richard R. Copley, Elisabeth Christians, Stefania Castagnetti

## Abstract

In eukaryotic cells, a spindle assembly checkpoint (SAC) ensures accurate chromosome segregation. This control mechanism monitors proper attachment of chromosomes to spindle microtubules and delays mitotic progression if connections are erroneous or absent. The SAC operates in all eukaryotic cells tested so far, but is thought to be relaxed during early embryonic development in animals. Here, we evaluate the checkpoint response to lack of kinetochore-spindle microtubule interactions in early embryos of diverse animal species from the main metazoan groups. Our analysis shows that there are two classes of embryos, either proficient or deficient for SAC activation during cleavage. Sea urchins, mussels and jellyfish embryos show a prolonged mitotic block in the absence of spindle microtubules from the first cleavage division, while ascidian and amphioxus embryos, like those of *Xenopus* and zebrafish, continue mitotic cycling without delay. SAC competence during early development shows no correlation with cell size, chromosome number or kinetochore to cell volume ratio, ruling out the hypothesis that lack of checkpoint activity in early embryos is due to the large egg volume. Our results instead indicate that there is no inherent incompatibility between SAC activity and large fast-dividing embryonic cells. We suggest that SAC proficiency is the default situation of metazoan embryos, and that SAC activity is specifically silenced in chordate species with fast dividing embryos.

## Introduction

The mitotic checkpoint, also known as the spindle assembly checkpoint (SAC), operates during mitosis and monitors bipolar attachment of spindle microtubules to kinetochores, specialized multi-protein complexes assembled on duplicated sister chromatids. In the absence of stable bipolar kinetochore-microtubule attachments the SAC generates an inhibitory signal, the mitotic checkpoint complex (MCC), which prevents activation of the anaphase-promoting complex/cyclosome (APC/C) and so delays chromosome segregation and mitotic exit. When all chromosomes have achieved bipolar attachments to microtubules, the SAC is quickly silenced resulting in APC/C activation, which leads to the proteolytic cleavage of securin and cyclin B1. Degradation of securin activates separase, thus resulting in cohesin cleavage and physical separation of sister chromatids, while cyclin B1 degradation inactivates cyclin dependent kinase (CDK), resulting in mitotic exit (Lara-Gonzalez et al., 2012; Musacchio 2015). This mechanism increases the fidelity of mitosis by preventing premature initiation of anaphase and subsequent generation of daughter cells with unequal chromosomal complements, a condition, known as aneuploidy, linked to cell and organismal lethality (Ricke and Deursen, 2013).

Despite the essential role of the SAC in achieving accurate chromosome segregation, genetic fidelity and reproductive success, microtubule perturbations that cause erroneous kinetochore-spindle associations do not trigger a robust spindle checkpoint response during the early rapid cell cycles (cleavage cycles) of embryonic development in fish and frog embryos. In *Xenopus laevis,* treatment with microtubule depolymerizing drugs does not delay the first 12 embryonic cycles and the associated oscillations of CDK activity, which continue with unchanged periodicity until the mid-blastula transition (MBT; Clute and Masui, 1995, Gerhart et al., 1984). Similarly, in zebrafish embryos, nocodazole treatment induces a metaphase arrest only after MBT (Ikegami et al., 1997, Zhang et al. 2015). In mouse, which like all mammals has slow (somatic-like) cleavage cycles compared to other animals, nocodazole treatment in 2-cell embryos causes a SAC-dependent mitotic delay (Kato and Tsunoda, 1992; Vázquez-Diez et al, 2019). These studies framed the hypothesis that the SAC is weak or silenced in early animal embryos especially those that undergo fast cleavage divisions. In such embryos, the SAC only becomes active later in embryogenesis, usually during early gastrulation, under the control of an as yet unidentified developmental timer (Clute and Masui, 1995; Zhang et al., 2015).

Recently an alternative hypothesis for the lack of a robust SAC response during early embryogenesis was brought to the fore by studies in *Caenorhabditis elegans* (Galli and Morgan 2016) that showed the ratio of kinetochore number to cell volume influences the strength of SAC response in embryos of these species. These findings are in keeping with earlier data from *Xenopus* egg extracts supplemented with high density sperm nuclei which displayed SAC activity at a kinetochore to volume ratio comparable to somatic cells (Minshull et al. 1994). Because a minimum signal threshold, dependent on the amount of Mad2 protein recruited on unattached kinetochores, needs to be reached to inhibit APC/C activity and elicit a SAC-mediated mitotic block (Collin et al. 2013), it was suggested that in large embryonic cells, the SAC is active but the signal generated by unattached kinetochores might be too dilute to trigger a significant checkpoint response (Galli and Morgan 2016). Thus, during early embryogenesis, the SAC would only become apparent when, following the decrease in cell size due to division without growth typical of embryonic cleavage, a sufficient kinetochore to cell volume ratio is reached. Contrary to this hypothesis, however, several earlier reports show that treatment with microtubule depolymerizing drugs delays cyclin B degradation and extends mitosis in zygotes of the sea urchins *Arbacia punctulata* and *Lytechinus variegatus* and of the clam *Spisula solidissima* (Sluder et al., 1994; Evans et al., 1983, Hunt et al., 1992), indirectly suggesting that the SAC may be effective in those embryos as early as the first cleavage, despite their large cell volume.

Here we use a comparative approach to assess the variability in SAC response during the early cell cycles of embryonic development in species representative of the main metazoan groups and to determine whether specific cellular characteristics, like cell size and kinetochore number, are good predictors of SAC competence. To complement the extensive data already available for vertebrates, we examined the mitotic response to complete microtubule depolymerization in early embryos of a range of invertebrate species. Unexpectedly, we found that lack of SAC activity is not a general feature of embryonic cleavage cycles. While ascidian (tunicate) and amphiouxus (cephalochordate) early embryos, like previously studied fish and frog embryos (vertebrates), continue to cycle without spindles, sea urchin and starfish (echinoderm), mussel (mollusk) and jellyfish (cnidarian) embryos show a prolonged checkpoint-dependent mitotic block from the first division in response to spindle perturbations. This species-specificity in SAC competence does not correlate with cell size, chromosome number or kinetochore to cell volume ratio, ruling out the hypothesis that SAC silencing during early development is due to the dilution of checkpoint signal in large cells. Instead our analysis suggests that silencing of SAC signaling during cleavage arose during animal evolution as a novel feature in the chordate lineage.

## Results

### Multispecies survey identifies two classes of embryos with different mitotic responses to spindle defects

The SAC monitors kinetochore-microtubule interactions and in somatic cells it delays mitotic progression in response to spindle defects. To assess SAC response during embryogenesis in diverse animal species, we monitored mitotic progression in the presence of the microtubule-depolymerizing drug nocodazole in 2-cell stage embryos from representative species of the main metazoan groups. To complement the extensive data already available in the literature for vertebrates (*Xenopus laevis* and *Danio rerio*) and nematodes (*Caenorhabditis elegans*), we chose the tunicate *Phallusia mammillata*, the echinoderms *Hacelia attenuata* (subphylum Asteroidea), *Paracentrotus lividus*, *Arbacia lixula*, *Sphaerechinus granularis* and *Strongylocentrotus purpuratus* (subphylum Echinoidea), the mollusk *Mytilus galloprovincialis* and the cnidarian *Clytia hemisphaerica*. In order to analyze SAC response under comparable conditions we used a concentration of nocodazole that completely depolymerized microtubules to generate a full set of unattached kinetochores. Treatment with 10 μM nocodazole after first cytokinesis (2-cell stage) completely depolymerized microtubules and blocked further cytokinesis in embryos of all selected species (S4 Figure). We then assessed mitotic progression by analyzing the mitotic marker Phospho Histone H3 (PH3; Hendzel et al, 1997) over the equivalent of at least one cell cycle time in control (+DMSO) and nocodazole (+ noco) treated embryos (Fig 1A). As shown in Fig 1B, we observed two qualitatively different responses to nocodazole treatment (red lines). In line with previous data from frog and fish, *P. mammillata* embryos continued to cycle in the presence of nocodazole, as evidenced by PH3 oscillation (Fig 1B*i*) occurring concomitantly with rounds of chromosome condensation and decondensation, suggesting lack of efficient SAC activation in these embryos. However, the response to microtubule depolymerization of embryos from all other analyzed species was strikingly different. Within 10-30 minutes of nocodazole treatment (depending on species), embryos showed condensed chromosomes and accumulated the mitotic marker PH3, indicating mitotic commitment. Both of these mitotic markers were maintained for the length of time equivalent to at least one cell cycle in the presence of nocodazole (Fig 1B*ii*–*viii*), and for some species, like the sea urchin *P. lividus* (Fig 1B*iii*), the mollusk *M. galloprovincialis* (Fig 1B*vii*) and the cnidarian *C. hemisphaerica* (Fig 1B*viii*), the mitotic arrest was extended up to two-to-three times their cell cycle length. Thus, contrary to the generally accepted dogma that early metazoan embryos lack spindle checkpoint activity, early embryos of echinoderm, mollusk and cnidarian species significantly delay mitotic progression in the presence of spindle defects.

**Figure 1:**
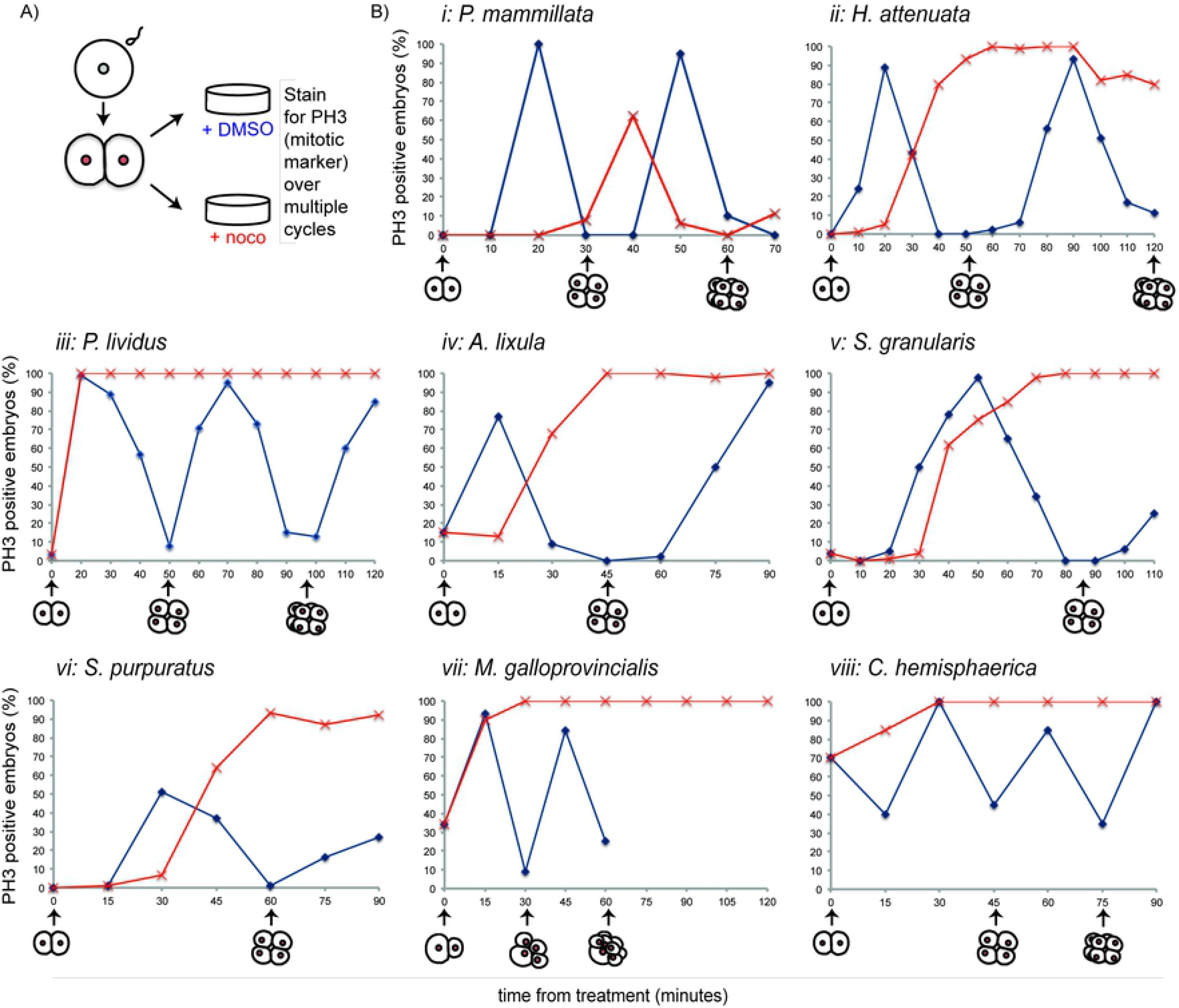
Nocodazole-induced spindle depolymerization defines two classes of embryos with qualitatively different mitotic responses. A) Schematic representation of assay used to test mitotic progression. Two-cell stage embryos of selected species were treated either with DMSO (0,1%) or 10 μM nocodazole and then fixed every 10 minutes for immunostaining with antibody against the mitotic marker PH3. B) Percentage of embryos accumulating PH3 in the presence of DMSO (blue) or 10 μM nocodazole (red) over the time of 2-3 cleavages (2 to 16 cell stage), representative of at least three independent experiments. 50-200 embryos were counted for each time point in each experiment. Drugs were added when 90% of embryos were at 2-cells (t0). For *M. galloprovincialis* control could not be quantified past 8 cells as divisions become asynchronous within each embryo.

### The mitotic delay observed in jellyfish, sea urchin and mussel embryos depends on the SAC kinase Mps1

To confirm that the mitotic delay observed in the presence of nocodazole is due to SAC activation, we further analyzed mitotic progression under conditions, which compromised SAC activity. In the presence of spindle defects, the SAC kinase Mps1 binds to unattached kinetochores where it regulates recruitment of other checkpoint components Bub1, BubR1, Bub3, Mad1 and Mad2 (Abrieu et al., 2001; Sacristan and Kops, 2015). In somatic cells, inhibition of Mps1 activity leads to displacement of SAC components from kinetochores, checkpoint inactivation and cell cycle resumption (Santaguida et al 2010; Abrieu et al. 2001). If the delay in mitosis observed in the presence of nocodazole is due to the activation of the SAC, then treatment with Mps1 inhibitors, like reversine (Santaguida et al., 2010), should restore mitotic timing in nocodazole treated embryos, resulting in mitotic exit and cell cycle resumption. For this analysis, we focused on a representative species from each animal group: *P. lividus* (echinoderm, Fig 2B-D), *C. hemisphaerica* (cnidarian, Fig 2E-G) and *M. galloprovincialis* (mollusk, Fig 2H-K). When embryos completed first cytokinesis, we treated them with 10 μM nocodazole alone or in combination with 0.5 μM reversine and assayed mitotic progression using several markers (Fig 2A). As already shown, in all three species, nocodazole alone caused an increase in mitotic index within 30 minutes of treatment, as evidenced by accumulation of cells with condensed chromosomes labeled with the mitotic marker PH3 (Fig 1B) and by the lack of nuclear membrane staining with the nuclear pore component Nup-153 (for *M. galloprovincialis,* Fig 2H). In nocodazole these mitotic markers were all maintained for at least the equivalent of two cell cycle times in all three species. Reversine treatment shortened the nocodazole-induced mitotic arrest, resulting in loss of chromatin associated PH3 staining (Fig 2B, E, J, K) and chromosome decondensation (Fig 2I, Hoechst). In mussel embryos, mitotic exit in reversine treated embryos was further confirmed by nuclear envelope reformation, as shown by Nup-153 staining (Fig 2H, I). Interestingly in all three species, PH3-labelled chromosomes started to accumulate again at later time points, indicating that cells that exited mitosis upon Mps1 inhibition then resumed the cell cycle and entered a new mitosis. Similar results were also obtained using another Mps1 inhibitor, AZ3146 (Hewitt et al., 2010), further validating that the release of the mitotic arrest observed in those embryos is due to specific inactivation of Mps1 activity (S5 Figure).

**Figure 2:**
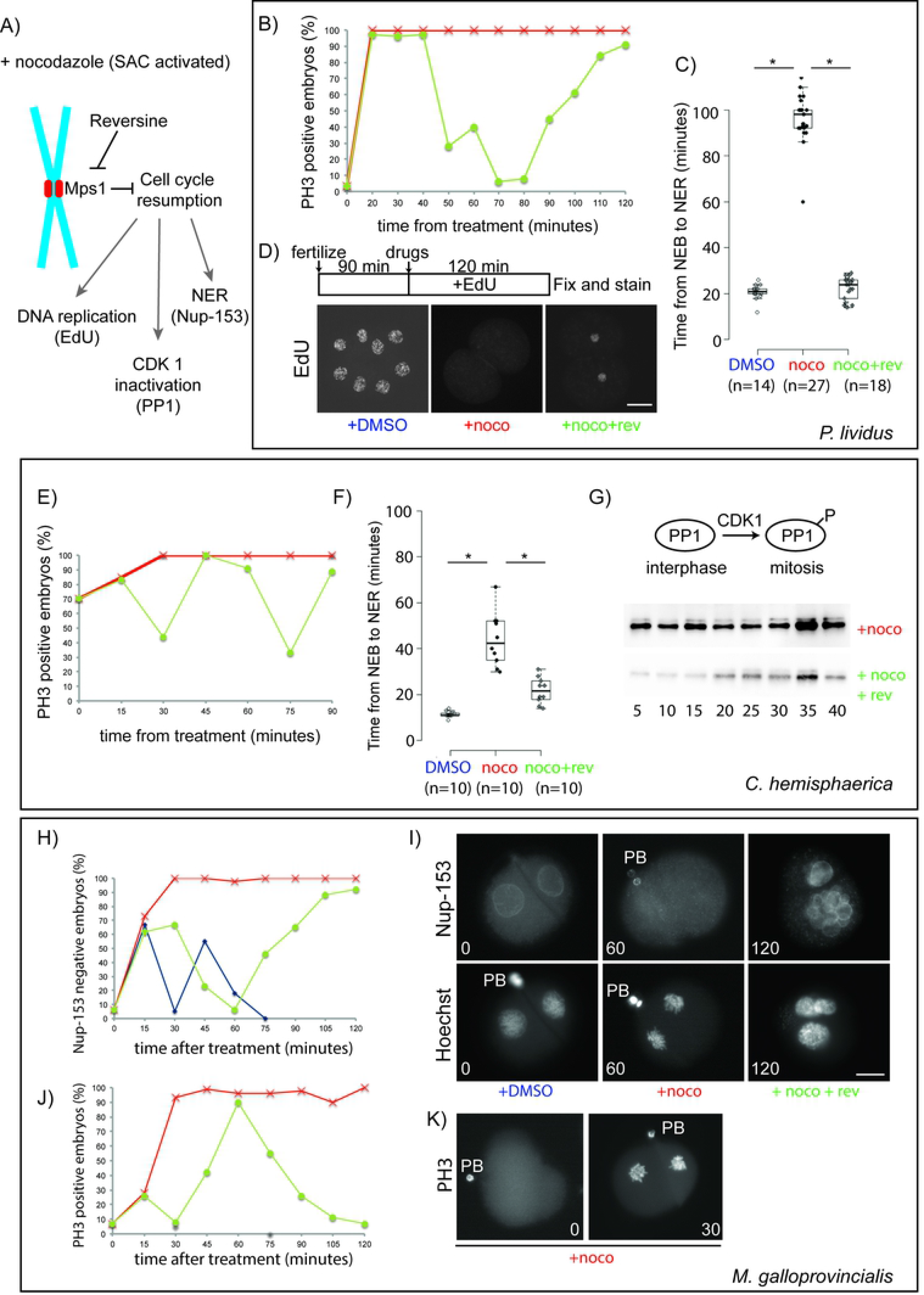
Microtubule depolymerization causes an Mps1-mediated mitotic block in cleavage stage embryos of *P. lividus*, *C. hemisphaerica* and *M. galloprovincialis*. A) Schematic representation of the effect of the Mps1 inhibitor reversine on cell cycle progression during SAC activation (+ nocodazole). B) Quantification of embryos accumulating PH3 in the presence of 10 μM nocodazole (red) or 10 μM nocodazole and 0.5 μM reversine (green) over time equivalent of two cell cycles. C) Quantification of length of mitosis in *P. lividus* embryos treated with 0.1% DMSO, 10 μM nocodazole or 10μM nocodazole and 0.5 μM reversine. Mitosis was measured as time from NEB to NER. Each dot represents one embryo. Boxes represent 25-75^th^ percentiles and the median is shown. Asterisks indicate statistical significance as determined by Student’s t-test, p<0.001. The numerical values associated with this graph are reported in S2 Table. D) Labeling of newly replicated DNA by EdU incorporation in control (+DMSO, left), nocodazole (middle) and nocodazole/reversine (right) treated embryos. EdU was added together with drugs (90 minutes post fertilization) when embryos reached 2-cell stage. All embryos were fixed when control reached 8-cell stage (210 minutes). 50 embryos were analyzed for each condition in 3 independent repeats. E) Quantification of PH3 positive *C. hemisphaerica* embryos in the presence of 10 μM nocodazole (red) or 10 μM nocodazole and 0.5 μM reversine (green). Representative of 4 independent experiments, n=20-30 for each time point. F) Quantification of length of mitosis in *C. hemisphaerica* embryos treated with DMSO, nocodazole or nocodazole and reversine. Each dot represents one embryo. Box plot parameters are as in C). G) Western blot analysis (5 embryos per lane) of phospho-PP1 in nocodazole (top) and nocodazole/reversine treated embryos (bottom). At 30 minutes, control embryos were at 4-cell stage. Timecourses in F and G come from independent experiments. H) Quantification of Nup153-labelled and J) PH3-labelled *M. galloprovincialis* embryos after treatment with 0.1% DMSO (blue), 10 μM nocodazole (red), or 10 μM nocodazole and 0.5 μM reversine (green). I) Representative images of embryos stained for Nup153 (top), DNA (Hoechst, bottom) and K) PH3. PB= polar body. Scale bar 30 μm.

For *C. hemisphaerica* and *P. lividus*, whose embryos are transparent, we could also measure the duration of mitosis in living embryos, as the time between nuclear envelope breakdown (NEB), which corresponds to prometaphase, and nuclear envelope reformation (NER), using DIC microscopy. In the presence of nocodazole, embryos of both species entered mitosis, as shown by the disappearance of discrete nuclei (NEB). NER, which marks exit from mitosis, was significantly delayed compared to control DMSO-treated embryos. In *P. lividus*, length of mitosis increased 5 fold, from 21 ±3 to 98 ±10 minutes, (Fig 2C), whereas in *C. hemisphaerica,* the interval between NEB and NER increased 3.5 fold, from 12 ±1 minutes in DMSO to 44 ±11 minutes in nocodazole (Fig 2F). Consistent with the results obtained with fixed embryos, inhibition of Mps1 activity, by reversine treatment, resulted in a significant reduction of the mitotic arrest observed in nocodazole-treated embryos (mitotic duration was shortened to 24 ± 5 minutes for *P. lividus* and to 22 ± 6 minutes for *C. hemisphaerica*, Fig 2C and 2F). Following mitotic exit reversine-treated embryos resumed cycling and re-entered mitosis as shown by subsequent rounds of NEB and NER. In *P. lividus* embryos, cell cycle resumption was further confirmed by visualization of DNA replication, using 5-ethynyl-2′-deoxyuridine incorporation (EdU). For this assay EdU was added to embryos in sea water at the time of drug treatment and embryos were fixed and stained after 2-cell cycles, when control embryos reached the 8-cell stage (Fig 2D). DNA replication, which was undetectable in nocodazole treated embryos over two cell cycle times (120 minutes), resumed following reversine treatment leading to nuclear staining in 2-cell arrested embryos within 80 minutes (Fig 2D). EdU incorporation was inefficient in *C. hemisphaerica* and *M. galloprovincialis* embryos and therefore this assay could not be carried out for these species.

Progression through the cell cycle is regulated by the activity of cyclin dependent kinases (CDKs) in combination with their activators, cyclins. Specifically, activation of the CDK1-cyclin B1 complex is required for mitotic commitment, and its inactivation drives mitotic exit. Cells arrested in mitosis have high CDK activity and phosphorylated mitotic targets, which are otherwise dephosphorylated upon mitotic exit. To confirm that reversine caused mitotic exit in *C. hemisphaerica*, we evaluated the phosphorylation status of PP1, a mitotic target of CDK-cyclinB1 (Wu et al, 2009, Lewis et al, 2013). As shown in Fig 2G, in the presence of nocodazole PP1 phosphorylation was maintained at a constant level for at least 60 minutes. Treatment with reversine caused a delay in mitotic entry. However once in mitosis, SAC impairment by reversine resulted in rapid loss of PP1 phosphorylation in nocodazole treated embryos, confirming that these embryos rapidly exited mitosis. Taken together these results show that in sea urchin, cnidarian and mollusk embryos, the mitotic block caused by spindle perturbations is SAC-dependent.

### SAC competence does not correlate with cell size across species

In our multispecies survey the tunicate *P. mammillata* was the only species that did not arrest in mitosis in the presence of nocodazole. We confirmed the lack of mitotic delay using live microscopy to follow nuclear behavior in DMSO and nocodazole treated embryos. Indeed both control and nocodazole treated embryos underwent multiple consecutive rounds of NEB and NER and chromosome condensation and decondensation (Fig 3A, nocodazole). Measurements of the duration of mitosis, as the time from NEB to NER, showed only a slight difference between DMSO and nocodazole treated embryos (<0.5 fold). In addition, as shown in Fig 3B we observed that as nocodazole-treated *P. mammillata* embryos underwent subsequent cell cycles, the length of mitosis remained unchanged, despite the increase in chromosome number and kinetochore to cell volume ratio, due to continuous cycling without intervening cytokinesis (Fig 3C). Consistent with previous observations in *C. elegans* (Galli and Morgan, 2016) and vertebrate tissue culture cells (Rieder and Cole, 2000), the length of interphase (I), measured as time from NER to NEB, instead increased at each cycle (Fig 3B). Thus, differently from echinoderm, cnidarian, nematode and mollusk early embryos, *P. mammillata* embryos lack SAC activity during embryonic cleavage.

**Figure 3:**
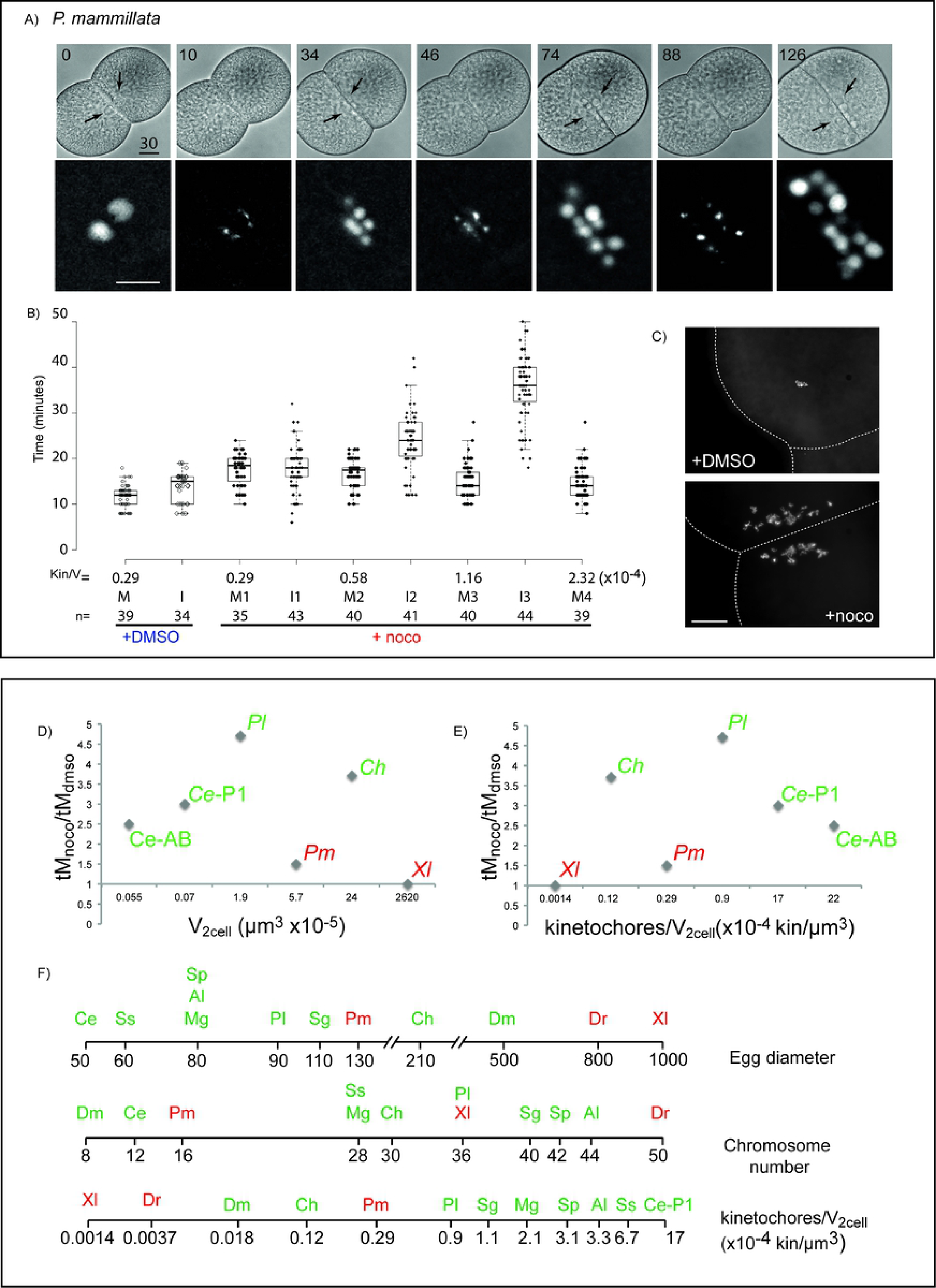
SAC response across species does not correlate with cell volume. A) Selected frames from a time-lapse movie of a *P. mammillata* embryo expressing the fluorescent DNA reporter, H2B-gfp, treated with 10 μM nocodazole after first cleavage. Numbers indicate time (minutes) from treatment. Arrows indicate nuclei visible in bright field optics. See also Movie 1. B) Length of mitosis (M, NEB to NER) and interphase (I, NER to NEB) in control (+DMSO), and nocodazole treated *P. mammillata* embryos. Kin/V indicates kinetochore to cell volume ratio following subsequent rounds of DNA replication. Box plots are as in Fig 2C. The numerical values associated with this graph are reported in S3 Table. C) Dapi stained chromosome spreads from control (DMSO) and nocodazole-treated (180 minutes) *P. mammillata* embryo. D,E) Ratio of average time spent in mitosis for nocodazole and DMSO treated embryos plotted against D) cell volume or E) kinetochore/ cell volume ratio in 2-cell stage embryos of different species. Ce= *C. elegans*, Pl= *P. lividus*, Pm= *P. mammillata*, Ch= *C. hemisphaerica* and Xl=*X. laevis*. For *Ce* as first division is asymmetric, volumes for both cells are presented (AB and P1). For *Ce* and *Xl* data were obtained from the literature. F) Egg diameter, chromosome number and kinetochore/cytoplasmic ratio at 2-cell stage for all species analyzed. For all species used in this study egg size was measured and is reported as average of 30-50 eggs. Red are species that do not delay mitosis, green are species that delay mitosis in the presence of nocodazole. Scale bar 30 μm.

As it was previously shown that the strength of SAC response can be modulated by cell size, we wondered whether the difference in mitotic response to spindle defects observed across species could be explained by the difference in cell size in early embryos, whose diameters range from tens of microns to millimeters depending on the animal species. We therefore compared cell size, kinetochore number and kinetochore to cell volume ratio at the 2-cell stage in all species used in our survey (Table 1). We included in this analysis also published data for *C. elegans* (Galli and Morgan, 2016), *X. laevis,* (Gerhart and Kirschner, 1984) and the fruit fly *Drosophila melanogaster* (Perez-Mongiovi et al., 2005). Comparison of the length of mitotic delay to egg size, chromosome number, cell volume or kinetochore to cell volume ratio at the 2-cell stage (Fig 3D, 3E and 3F) showed that the difference in SAC response across species does not correlate with any of these parameters; in fact large cells with low kinetochore to cell volume ratio, like those of 2-cell *C. hemisphaerica* embryos delay mitosis more efficiently than the cells of smaller embryos, like *C. elegans* or *P. mammillata*. In addition, by the 4^th^ mitotic cycle, *P. mammillata* nocodazole treated embryos reach the same kinetochore to cell volume ratio of SAC proficient *P. lividus* 2-cell embryog, but do not delay mitotic progression (Fig 3B).

**Table 1.** Morphometric data for embryos of all analyzed species.

| **Table 1: Morphometric data for embryos of all analyzed species** |  |  |  |  |  |  |  |
| --- | --- | --- | --- | --- | --- | --- | --- |
|  | Species | Oocyte diameter (μm) | Chromosome number (2n) | Kin/V<sub>2cell</sub> ×10<sup>-4</sup> (kin/μm<sup>3</sup>) | V<sub>nucleus</sub> (μm<sup>3</sup>) | Spindle length (μm) | SAC competence |
| Chordates | *Xenopus laevis* | 1000 | 36 | 0,0014 | 3053<sup>a</sup> | 53,5<sup>b</sup> | –<br>Gerhart et al. 1984 |
|  | *Danio rerio* | 800 | 50 | 0,0037 |  |  | –<br>Ikegami et al. 1997 |
|  | *Ciona intestinalis* | 140 | 28 | 0,39 |  |  | –<br>this study |
|  | *Phallusia mammillata* | 130 | 16 | 0,29 | 1150 | 20-30 | –<br>this study |
|  | *Branchiostoma lanceolatum* | 130 | 38 | 0,66 |  |  | –<br>this study |
| Echinoderms | *Hacelia attenuata* | 240 |  |  |  |  | +<br>this study |
|  | *Paracentrotus lividus* | 90 | 36 | 0,9 | 1437 | 18<sup>c</sup> | +<br>this study |
|  | *Arbacia lixula* | 80 | 44 | 3,3 |  |  | +<br>this study |
|  | *Strongylocentrotus purpuratus* | 80 | 42 | 3,1 |  |  | +<br>this study |
|  | *Sphaerechinus granularis* | 110 | 40 | 1,1 |  |  | +<br>this study |
| Mollusks | *Mytilus galloprovincialis* | 80 | 28 | 2,1 | 2352 |  | +<br>this study |
|  | *Spisula solidissima* | 60 | 28 | 6,7 |  |  | +<br>Evans et al. 1983 |
| Arthropods | *Drosophila melanogaster* | 500 | 8 | 0,018 |  |  | +<br>Perez-Mongiovi et al. 2005 |
| Nematodes | *Caenorhabditis elegans* | 50 | 12 | AB= 22<sup>f</sup><br>P1= 17<sup>f</sup> | 523<sup>d</sup> | 13<sup>e</sup> | +<br>Encalada et al. 2005 |
| Cnidarians | *Clytia hemisphaerica* | 210 | 30 | 0,12 | 526 | 30-35 | +<br>this study |

<sup>a</sup> Levy DL and Heald R. 2010. Nuclear size is regulated by importin $\alpha$ and Ntf2 in *Xenopus*. *Cell* 143:288-298
<sup>b</sup> Wühr M, Chen Y, Dumont S, Groen AC, Needleman DJ, Salic A and Mitchison TJ. 2008. Evidence for an upper limit to mitotic spindle length. *Curr Biol* 18:1256-61
<sup>c</sup> Lacroix B, Letort G, Pitay L, .... and Dumont J. 2018. Microtubule dynamics scale with cell size to set spindle length and assembly timing. *Dev Cell* 45:496-511
<sup>d</sup> Ladouceur AM, Dorn JF and Maddox PS. 2015. Mitotic chromosome length scales in response to both cell and nuclear size. *J Cell Biol* 209:645-51
<sup>e</sup> Greenan G, Brangwynne CP, Jaesch S, Gharakhani J, Jülicher F and Hyman AA. 2010. Centrosome size sets mitotic spindle length in *Caenorhabditis elegans* embryos. *Curr Biol* 20:353-358
<sup>f</sup> Galli M and Morgan DO. 2016. Cell size determines the strength of the spindle assembly checkpoint during embryonic development. *Dev Cell* 36:344-352

Since nuclear and spindle size also vary among different cell types and during development we also checked whether SAC competence could be related to changes in either of these features. Although we could not obtain measurements for all species analyzed in our survey, we observed no correlation with either of these two parameters. For example, SAC-competent *P. lividus* and SAC-deficient *P. mammillata* blastomeres have comparable sized nuclei (14.4 ± 1μm and 13 ± 2μm, respectively) and both are smaller than that of *M. galloprovincialis,* 16,5 ± 2μm (Table 1). Similarly, SAC deficient *P. mammillata* blastomeres have spindles of intermediate size between SAC proficient *P. lividus* and *C. hemisphaerica* (Table 1).

Thus, our data show that cell, nuclear and spindle size, chromosome number and kinetochore to cell volume ratio are not good predictors of SAC activity during early embryonic development.

### Chordate embryos do not arrest in mitosis in the presence of spindle perturbations

In the multispecies analysis shown above only the tunicate (*P. mammillata)* and vertebrate (*X. laevis* and *D. rerio*) embryos failed to trigger a mitotic delay in response to spindle defects during cleavage. Because tunicates and vertebrates, together with cephalochordates, form the chordate clade (Fig 4A), we asked whether lack of SAC activity during cleavage is a common feature of chordate embryos (excluding mammals, which have highly atypical early development featuring slow, somatic-type cell cycles). To address this question, we analyzed the mitotic response to microtubule depolymerization in *Ciona intestinalis,* another tunicate species, and in *Branchiostoma lanceolatum* a species representative of the cephalochordate group.

**Figure 4:**
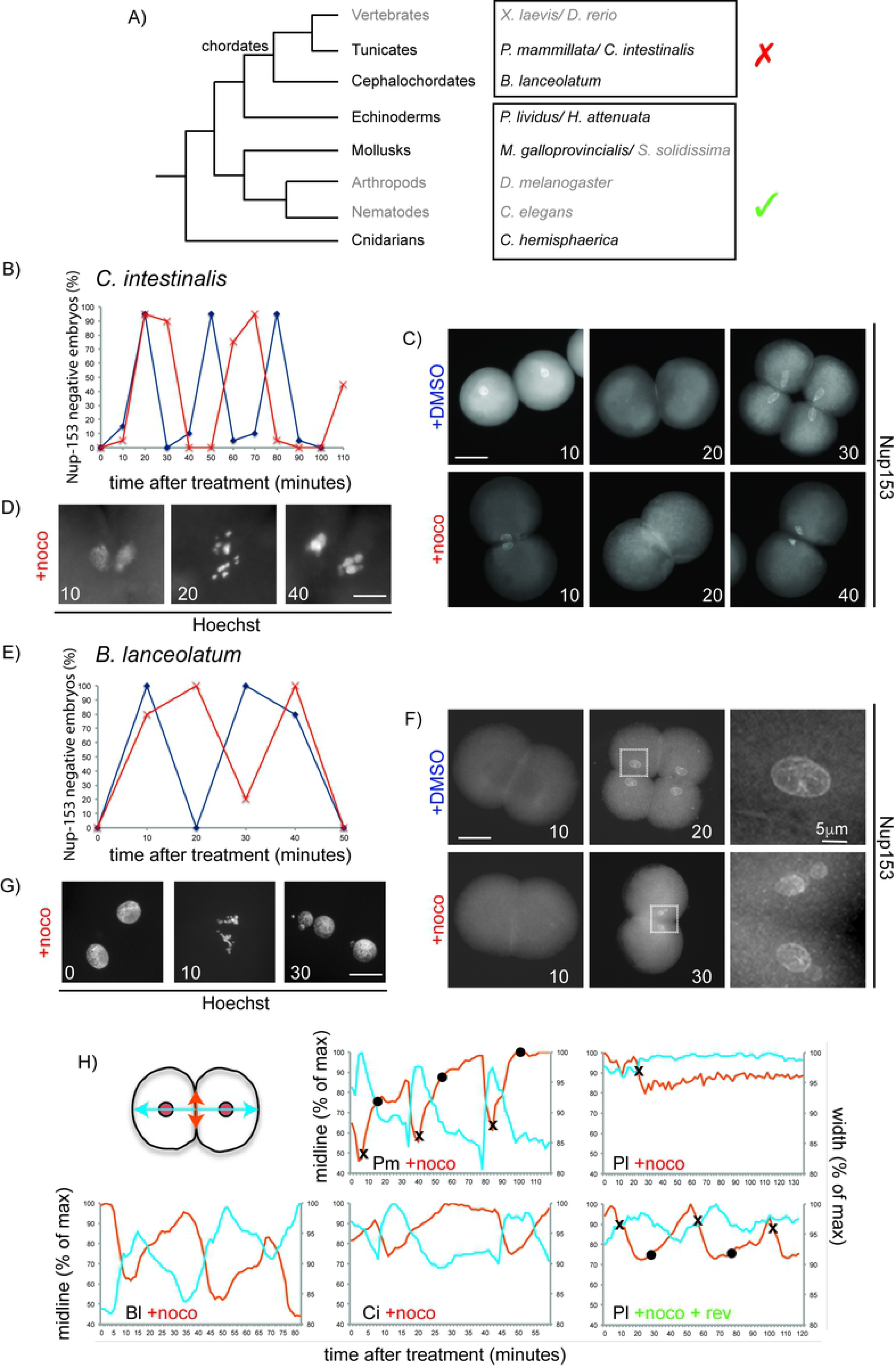
Nocodazole treatment does not delay mitotic progression during cleavage in chordate embryos. A) Phylogenetic tree indicating phyla, analyzed species and their SAC response. Species analyzed in this study are in black, species for which information has been obtained from the literature are in grey. B) Quantification of Nup153-negative *C. intestinalis* embryos (without nuclei=in mitosis) in the presence of DMSO (blue) or 10 μM nocodazole (red) over the time of 3 divisions (2-16 cells). Representative of 3 independent experiments. n=20-30 embryos per time point for each repeat. C) Representative Nup-153-stained embryos and D) Hoechst stained nuclei for *C. intestinalis*. E) Percentage of *B. lanceolatum* embryos without nuclei, as determined by Nup-153 staining, in the presence of DMSO (blue) or 10 μM nocodazole (red) over the time of two divisions (2-8 cells). Representative of 3 independent experiments. n=50-100 embryos per time point. F) Representative Nup-153 and G) Hoechst stained nuclei for *B. lanceolatum*. Duration of treatment is indicated on each image (in minutes). H) Measurement of long axis of embryo (width, blue) and cell-cell contact region (midline, orange) during 2-3 cell cycles in a representative embryo of *B. lanceolatum* (Bl), *P. mammillata* (Pm, see movie 1), *C. intestinalis* (Ci) and *P. lividus* (Pl) in the presence of 10 μM nocodazole (+noco) or for Pl 10 μM nocodazole and 0.5 μM reversine (Pl +noco+rev). Measurements are reported as percentage of maximum length throughout the recording. Crosses (✖) correspond to NEB, and circle (•) to NER. Scale bar 30 μm.

As shown in Fig 4, the response of both species to nocodazole treatment was very similar to vertebrates and *P. mammillata.* Although nocodazole treatment blocked cytokinesis, nuclei of both *B. lanceolatum* and *C. intestinalis* embryos continued to cycle and underwent several rounds of mitosis, as evidenced by subsequent rounds of chromosome condensation and decondensation (Fig 4D and 4G), and of nuclear envelope breakdown and reformation (Fig 4B and 4C for *B. lanceolatum* and Fig 4E and 4F for *C. intestinalis*). As neither of these embryos is transparent, nuclear dynamics could not be followed in vivo. However, time-lapse microscopy revealed that these nocodazole-treated embryos underwent cyclic shape changes, whereas the SAC arrested *P. lividus* embryo did not. As most animal cells acquire a round shape upon mitotic entry (Lancaster et al., 2013), we used shape change as a marker of progression through the cell cycle. We quantitated embryo shape over time, by imaging embryos in the presence of nocodazole for at least 2 complete cell cycles and then measuring two parameters (Fig 4H): contact region between the two blastomeres (midline, in orange) and total width of the embryo (long axis, in blue). In *P. mammillata, C. intestinalis* and *B. lanceolatum* nocodazole-treated embryos, midline length and embryo width oscillated cyclically and in a reciprocal fashion. This resembles the periodic rounding and flattening documented for *X. laevis* eggs induced to cycle in the absence of cell division (Hara et al., 1980). For *P. mammillata*, whose nuclei are easily visible, NEB was observed when the midline was at its shortest and the embryo width at its longest, and NER took place once cells regained full adhesion (longest midline, shortest width). In contrast SAC proficient *P. lividus* embryos, midline length and embryo width remained essentially constant throughout the nocodazole-induced mitotic arrest (Fig 4H, Pl +noco). When SAC signaling was inhibited by reversine treatment, cyclic cell shape changes resumed with mitosis (NEB-NER) corresponding to periods of minimal blastomere contact (Fig 4H, Pl +noco+rev). Taken together these results show that *C. intestinalis* and *B. lanceolatum*, like *P. mammillata*, fish and frog embryos, continue to cycle in the presence of spindle perturbations and are therefore not SAC competent during early embryonic development. Thus, silencing of the SAC during cleavage may be associated with the emergence of the chordate lineage during animal evolution.

## Discussion

### SAC activity in embryos defines two classes of animals

The spindle assembly checkpoint operates during mitosis to delay the onset of anaphase under conditions that could otherwise compromise accurate chromosome segregation (Musacchio and Salmon 2007), and is thus important for cell and organismal viability. Despite this essential function, it has long been thought that the SAC is inefficient in early development of animal embryos with large eggs, as they undergo fast cycles. Here, we have undertaken a rigorous survey of the SAC response to spindle defects in embryos of diverse animal species, combining both new experimental data and previous findings from the literature. Because different microtubule poisons can provoke variable levels of SAC activity (Collin et al., 2013), we included in our analysis only studies performed using the microtubule depolymerizing drug nocodazole at a concentration that completely depolymerizes spindle microtubules, therefore generating a full complement of unattached kinetochores and maximum SAC signal. Our analysis shows that in the presence of unattached kinetochores, mitotic progression is unperturbed in fish, frog, amphioxus and ascidian embryos, whereas sea urchin, mussel, jellyfish, nematode and insect embryos delay mitotic exit. We conclude that there is no inherent incompatibility between the fast division typical of cleavage-stage embryonic development and spindle checkpoint activation.

### SAC activity in relation to kinetochore and cytoplasm content

Variations in the length of mitotic delay induced by SAC activation have been reported previously in several cellular contexts and were partially attributed to differences in cell size and kinetochore to cell volume ratio (Mishull et al., 1994; Galli and Morgan, 2016, Kyogoku and Kitjima, 2017). However, our analysis shows that the presence of SAC competence during embryo cleavage cycles does not correlate with reduced cell size across different species, with large jellyfish (diameter 210 μm) and starfish (240 μm) embryos mounting a prolonged block from first division and the smaller ascidian (130-140 μm) and amphioxus embryos (130 μm) not delaying mitosis for several divisions (Table 1). Likewise, pairwise comparisons also suggest that chromosome number (*P. lividus* and *X. laevis:* 36 chromosomes*; M. galloprovincialis* and *C. intestinalis:* 28 chromosomes) and kinetochore to cell volume ratio (*P. lividus* and *B. lanceolatum*) are not strong indicators of SAC competence at the egg-to-embryo transition across metazoans (Table 1). Consistent with this conclusion, it was previously reported that in *D. melanogaster*, whose eggs are 500 μm long and have only 4 chromosome pairs, treatment of stage 3-6 syncytial embryos with colchicine arrests the nuclear cycle at a prometaphase-like stage, suggesting that the SAC is active from early cleavage stage in these large insect cells (Perez-Mongiovi et al. 2005, Sullivan et al. 1993). However, in early *Drosophila* embryos cyclin B degradation and CDK inactivation occur only locally (Su et al. 1998) in the area of the spindle rather than at the level of the whole embryo. This observation raises the possibility that the SAC may be regulated locally in the vicinity of the chromosomes. Indeed, previous work carried out in PtK_1_ cells containing two separate spindles showed that once all kinetochores are attached to spindle microtubules within one spindle, anaphase will start irrespective of the presence of unattached kinetochores on the second spindle, suggesting that SAC signal is not diffusible (Rieder et al 1997). If SAC action is limited to the spindle region then the strength of the SAC response may be a function of the volume of a subcellular region local to the spindle area, rather than total cell volume. Spindle size itself, defined as pole to pole distance in metaphase, however, was shown to scale linearly with cell size across embryos of many different species (Crowder et al. 2015) and we confirmed this trend for our species. Thus for 2-cell embryos, difference in spindle size is unlikely to explain the difference in SAC activity observed across same sized embryos. We can conclude that there is no straightforward link between any of the cellular parameters that we analyzed, which include kinetochore number, spindle length, volume of cytoplasm or nucleus and the categorization of species into SAC proficient and SAC deficient embryos.

### SAC deficient embryos as an evolutionary novelty in the chordate linage

While we could not uncover any physical explanation for variation in SAC efficiency, it was immediately apparent when looking at their phylogenetic grouping that all species with SAC-deficient embryos are chordates, while species in all non-chordate clades possess SAC-competent embryos (Fig 4A). Thus loss of the SAC in cleaving embryos may be associated with the emergence of the chordate lineage. Sampling a wider number of species and metazoan groups under these same experimental conditions will be required to test this hypothesis further. Some supporting examples of non-chordate species being SAC positive can already be inferred from the literature, although the use of different drugs and assays to assess mitotic progression complicates comparisons. As already mentioned, colchicine treatment blocks cyclin B degradation in clam embryos during first mitosis (Hunt et al, 1992), and addition of nocodazole blocks nuclear division in embryos of another mollusk, the gastropod *Ilyanassa obsoleta* (Cather et al. 1986). Similarly, treatment with colchicine delays nuclear division at least for the length of one cell cycle in embryos of the fruitfly *D. melanogaster* (Perez-Mongiovi et al, 2005) and in binucleated embryos of the gall midges *Wachtliella periscariae* (Wolf 1978) and *Heteropeza pygmaea* (Kaiser and Went 1987).

Combining all available data, we can propose that SAC proficiency during sexual reproduction is an ancestral feature of metazoan embryos, and that SAC signaling became silenced during early development in chordate embryos. At the mechanistic level, lack of SAC activity in chordates could simply reflect absence in the egg of one or more of the basic SAC components. Although further analysis will be required, we do not favor this possibility since in our analysis of available transcriptomic data, we have determined that Mad1, Mad2, Bub1, Bub3 and Mps1 (S1 Table) are expressed at the mRNA level both before and after fertilization in species which have no SAC activity during cleavage, like *P. mammillata*, *C. intestinalis* (Aniseed, Brozovic et al., 2018) and *B. lanceolatum* (Oulion et al., 2012; Marletaz et al., 2018, and H. Escriva personal communication). Moreover checkpoint proteins, like XMad1 and XMad2, are present in the cytoplasm of SAC-deficient *X. laevis* early embryos (Chen et al. 1998). A number of scenarios can be envisaged to explain the lack of SAC activity in the presence of SAC components. Kinetochores may be modified to hinder their recognition by the checkpoint machinery or to interfere with the efficiency of MCC generation. Alternatively, as already suggested for mouse embryos, changes in the relative concentrations of SAC components and APC/C may result in an imbalance between inhibitor and target, effectively silencing checkpoint function. Finally an as yet unidentified SAC inhibitor may be present in chordate eggs and embryos and function to silence spindle checkpoint signaling during early development.

Given our current understanding of SAC function in maintaining ploidy, it is hard to understand what selective advantage could be associated with loss of SAC signaling in chordate embryos. At this point we can only speculate that SAC silencing is a by-product of some other change in reproductive regulation or oogenesis that could impact the levels of mitotic molecular regulators or the availability of kinetochores. The one exception to chordate SAC deficiency concerns mammalian embryos, which have undergone an extreme shift in reproductive strategy to viviparity, allowing the cleaving embryo to reduce its dependence on oocyte nutrient reserves and to lengthen their cell cycle and the duration of mitosis. In both mouse and human embryos, however, the SAC is highly inefficient leading to the formation of mosaic-aneuploid embryos (Bolton et al., 2016; Vanneste et al., 2009). Notably, in mouse pre-implantation embryos the presence of several unattached kinetochores fails to prevent mitotic progression, but extending the duration of mitosis improves SAC efficiency and reduces chromosome segregation errors (Vázquez-Diez et al., 2019), supporting a possible relationship between lengthening of mitosis and acquisition of SAC activity in mammalian embryos. Further analyses will be required to understand the underlying molecular mechanism controlling spindle checkpoint control during early development in these classes of embryos and the possible links between these changes in mitotic control and evolutionary transitions.

## Materials and methods

### Gamete collection and fertilization

*P. lividus*, *A. lixula*, *S. granularis* and *H. attenuata* adults were collected from the bay of Villefranche-sur-mer (France), *P. mammillata* and *M. galloprovincialis* at Sète (France), *C. intestinalis* at Roscoff (France) and *B. lanceolatum* at Argelès-sur-Mer (France). All these species were maintained in aquaria by CRBM at the Laboratoire de Biologie du Developpement de Villefranche-sur-mer (LBDV). *S. purpuratus* adults were obtained from Patrick Leahy (Kerchoff Marine Laboratory, California Institute of Technology, Pasadena, CA, USA) and kept in aquaria at University College London (UCL, London UK).

*S. purpuratus* adults were induced to spawn by injection of 0.55 M KCl and all manipulations were carried out at 15°C. For the other three sea urchin species, gametes were obtained by dissection and all manipulations were carried out at 18-20°C; eggs were collected in microfiltered sea water (MFSW) and used within the day, whereas sperm was collected dry and maintained at 4°C for up to a week. Prior to fertilization eggs were filtered to remove ovarian tissue and debris (100 μm filter pore size for *P. lividus* and *S. granularis*, 70μm for *A. lixula*). When removal of the fertilization membrane was required (for immunofluorescence) eggs were treated with 1X FC (10 μM 3-amino-1,2,4-triazole, 5 μM EDTA, 200 μM Tris-HCl pH8.2) for 2-3 minutes prior to fertilization to prevent hardening of the membrane. The fertilization membrane was removed by filtration (70μm for *P. lividus* and *S. granularis* and 54 μm for *A. lixula*) and excess sperm was removed by rinsing twice in MFSW.

For *H. attenuata* gametes were obtained by aspiration through the arm using a syringe with 18G needle. Oocytes were immediately matured with 10μM 1-methyladenine (1-MA, Sigma-Alderich) and after 13 minutes they were fertilized in glass dishes and cultured at 21°C.

For *P. mammillata* and *C. intestinalis* gametes were obtained by dissection. Dry sperm was maintained at 4°C, and eggs were dechorionated in 0.1% trypsin for *P. mammillata* or in pronase/thioglycolate for *C. intestinalis* as described (Sardet et al., 2011). All manipulations were performed at 18°C in dishes coated with gelatin or agarose to prevent adhesion and lysis (Sardet et al., 2011). Prior to fertilization sperm was activated by resuspension in basic seawater (pH 9.2) for 20 minutes.

*B.lanceolatum* mature adults were maintained at 16-17°C and induced to spawn by thermal shock at 23°C for 36 hours, as previously described (Theodosiou et al., 2011). Oocytes were collected in petri dishes and fertilized with a dilution of fresh sperm, and developing zygotes were incubated in MFSW at 19°C (Thedosiou et al., 2011).

*C.hemisphaerica* eggs and sperm were obtained by light induced spawning from animals raised in the laboratory and maintained at 19°C in artificial sea water (Houliston et al., 2010).

*M. galloprovincialis* adults were maintained in sea water at 15°C. To induce spawning animals were transferred into individual containers with sea water at 24°C, after rigorous cleaning and brushing of animal shells. Oocytes were fertilized in petri dishes and embryos developed at 17°C.

### Drug treatments

All drugs were maintained as stock solutions in DMSO at −20°C and diluted as appropriate in MFSW prior to usage. Nocodazole (Sigma, 33 mM stock solution in DMSO) was used at a final concentration of 10 μM, reversine (Axon Medchem, 5mM stock solution in DMSO) was used at a final concentration of 0.5 μM and AZ3146 (Santa Cruz Biotechnology, stock solution 22 mM in DMSO) was used at a final concentration of 2 μM.

In all experiments drugs were added when 90-95% of the embryos reached 2-cell stage to avoid regression of the cleavage furrow, and drug treatment was then maintained for the entire length of the experiment. Each experiment was repeated between 3-5 times.

### Immunofluorescence

For immunofluorescence, embryos were fixed overnight in −20°C 90% methanol containing 50 mM EGTA. After fixation embryos were washed 3 times in PBS containing 0.1% tween 20, then preblocked in PBS containing 3% BSA for 1 hour at room temperature and then incubated overnight at 4°C in PBS containing 3% BSA and the appropriate dilution of primary antibody. The mouse anti-PH3 (phospho S10, Abcam) antibody was diluted 1:1000, the rabbit anti-PCNA (Sigma-Alderich) 1:100, the mouse anti-Nup153 (Covance) 1:500. Following 3 washes in PBS-0.1% Tween20, embryos were incubated with specific fluorescently-labelled secondary antibodies at room temperature for 1-2 hours. Following 2 further washes in PBS-0.1% Tween20, embryos were incubated for 10 minutes in PBS+ 0.1% Tween20 containing Hoechst (5μg/ml), washed twice and then mounted in citifluor AF1 (Science Services) for imaging and quantification. Each experiment was repeated 3-5 times and depending on the species 25-200 embryos were counted for each sample.

### EdU staining

EdU staining was performed using the Click-iT EdU Imaging kit (Invitrogen), following the protocol provided by the manufacturer. Briefly, 10μM EdU was added to 2-cell stage *P. lividus* embryos in MFSW at the same time as DMSO or drugs. Embryos were maintained in EdU for 1 to 3 generation times and then fixed in 3.7% paraformaldehyde/PBS for 15 minutes at room temperature. Following 2 washes in PBS containing 0.1% TritonX100, embryos were permeabilized in PBS-0.5% TritonX100 for 20 minutes at room temperature and washed again twice in PBS containing 3% BSA. Following a 30 minute click-IT labelling reaction, embryos were washed extensively in PBS-0.1% TritonX100 and mounted in citifluor AF1 for imaging.

### Western blots

To prepare protein extracts of *C. hemisphaerica,* 5 embryos were lysed in Laemmli sample buffer (50 mM Tris-HCl (pH 6.8), 2% SDS, 0.1 % bromophenol blue, 10 % glycerol, 100 mM dithiothreitol) at 5 minute intervals starting from the 2-cell stage. Protein samples were separated on 10% SDS-polyacrylamide gels and transferred to nitrocellulose membranes. After blocking in 3% BSA, to preserve phospho-antigens, membranes were incubated over night at 4°C with mouse anti-phosho-PP1 antibody (Wu et al., 2009) (pospho-T320 Abcam, 1:1000) After washing, membranes were incubated with anti rabbit horseradish peroxidase-conjugated secondary antibody (Jackson ImmunoResearch 1:10000) and detection was carried out with SuperSignal West Pico chemiluminescent substrate (Thermo Scientific) as described by the manufacturer.

### Chromosome spreads

*P. mammillata* embryos were treated with DMSO or 10 μM nocodazole for 120 minutes, then washed in hypotonic solution (75 mM KCl), then in 37.5 mM KCl and finally washed four times in cold methanol:acetic acid (3:1), before fixation at −20°C overnight in methanol:acetic acid (3:1). After washing in 60% acetic acid, a few droplets of acetic acid containing the embryos were dripped onto cold methanol-washed slides from about 20 cm hight, air dried, and mounted in 50% glycerol containing DAPI for imaging with a Leica SP5 confocal microscope.

### Time-lapse microscopy and microinjection

Two cell stage embryos of *P. mammillata, C. intestinalis, B. lanceolatum, or P. lividus* were placed in sea water containing appropriate drugs in glass bottom dishes (MatTek corporation) or mounted between gelatin-coated slide and coverslip using Dow Corning vacuum grease as spacer as described (Sardet et al., 2011) and filmed on a Zeiss Axiovert inverted microscope using bright field optics, 20X or 40X objective lenses, and Metamorph acquisition software. To observe chromatin dynamics, *P. mammillata* eggs were injected before fertilization with synthetic mRNA encoding histone H2B fused to GFP (see McDougall et al., 2015 for construct and methods).

## Acknowledgments

We are grateful to L. Gilletta for animal care, to E. Houliston for discussion and critical reading of the manuscript, to P. Oliveri, R. Dumollard, C. Hebras, G. Pruliere, V. Costache, S. Chevalier, T. Momose, L. Leclère, H. Yasuo, G. Williaume, F. Lahaye, M. Schubert, E. Zieger, and H. Escriva for assistance, discussion and sharing of animals and unpublished data. This work was supported by funding from the CNRS and from PACA region (project Mepanep, 2014_03738). The authors declare no competing financial interests.

## List of supporting information

S1_Table.pdf Sequences of analyzed SAC components

S2_Table.xlsx Numerical values associated with Figure 2C

S3_Table.xlsx Numerical values associated with Figure 3A

S4_Figure.tiff Nocodazole treatment eliminates microtubules

S5_Figure.tiff The Mps1 inhibitor AZ3146 releases the nocodazole-induced mitotic block observed in *P. lividus* and *C. hemisphaerica*

Movie 1: *P. mammillata* 2-cell embryos expressing H3B-Venus, in the presence of 10μM nocodazole.

